# Adaptive optics enhanced all-optical photoacoustic tomography

**DOI:** 10.1101/2021.01.13.426260

**Authors:** Jakub Czuchnowski, Robert Prevedel

## Abstract

All-optical ultrasound detection bears a number of unique advantages for photoacoustic tomography, including the ability for high resolution sampling of the acoustic field and its compatibility with a wide variety of other optical modalities. However, optical schemes based on miniaturized cavities are sensitive to optical aberrations as well as manufacturing-induced cavity imperfections which degrade sensor sensitivity and deteriorate photoacoustic image quality. Here we present an experimental method based on adaptive optics that is capable of enhancing the overall sensitivity of Fabry-Pérot based photoacoustic sensors. We experimentally observe clear improvements in photoacoustic signal detection as well as overall image quality after photoacoustic tomography reconstructions when applied to mammalian tissues *in vivo*.

## 1. Introduction

Photoacoustic tomography is a non-invasive deep-tissue imaging modality that combines optical contrast with high-resolution ultrasound detection to enable high resolution imaging in deep, scattering tissues [1]. Multiple detector types and geometries were developed over the years [1], with optical methods for acoustic wave detection gaining increasing attention (see [2] for review). Here, the use of planar Fabry-Pérot (FP) cavity sensors has been particularly promising, as it combines high sensitivity with the ability to measure acoustic waves at well-defined spatial locations (given by the interrogating beam size on the sensor), which is important for high-resolution tomographic image reconstruction [3, 4]. In this approach, a pressure sensitive FP cavity is formed by sandwiching a layer of elastomere (e.g. Parylene C) between two dichroic mirrors. This allows the cavity to deform elastically upon incidence of a pressure wave, thus modulating the position of the FP interferometer’s transfer function (ITF) which depends on the (optical) thickness of the cavity. By tuning the interrogation laser wavelength to the point of maximum slope on the ITF (so-called bias wavelength) one can obtain maximum sensitivity to spatial displacements, which in turn enables to optically detect and maximally amplify the incident acoustic wave. Experimentally, this approach has enabled acoustic sensing in the range of 100*−*10^6^ Pa with a broadband frequency response (bandwidths up to 40*∼*MHz) [3, 4]. At the same time, the sensors’ dichroic mirrors can be designed appropriately to allow for efficient delivery of excitation light or the combination with other optical imaging modalities.

As light based sensors, FP interferometers (FPIs) are sensitive to both beam and cavity aberrations which can limit their performance under certain practical conditions. FP cavities are especially sensitive to wavefront aberrations as their optical performance requires high spatial uniformity of the light beam for efficient interference [5]. Recently, two theoretical frameworks were developed that enable to study the effects of both beam and cavity aberrations on the overall sensitivity performance [6, 7]. Furthermore, proof-of-principle demonstrations also showed that this loss can be partially compensated by the use of optical aberration correction approaches based on Adaptive Optics (AO) ([6], **Figure 1a**). This previous work, however, focused on theoretical aspects of the light-cavity interaction and their experimental validation was limited to a point-wise characterisation of the interferometer’s sensitivity. In particular, it did not address the question to which extent AO can be utilized in practical settings to improve the detected PA signal amplitude and image quality. In this paper, we investigate the experimental requirements for AO-enhanced photoacoustic tomography under realistic imaging conditions. Specifically, we show how focal spot shifts, induced by the AO wavefront correction, can be experimentally tracked and actively corrected. This is crucial for a spatially accurate sampling of the acoustic field over the entire FPI active area which is important for large-scale and high-resolution 3D image reconstruction as well as to ensure optimal convergence of the iterative AO routine. Using our developed routine, we show that readily available, pre-calibrated deformable mirrors are sufficient to achieve significant improvements in optical sensitivity and photacoustic signal level which ultimately translate to improvements in PA image quality.

**Figure 1:**
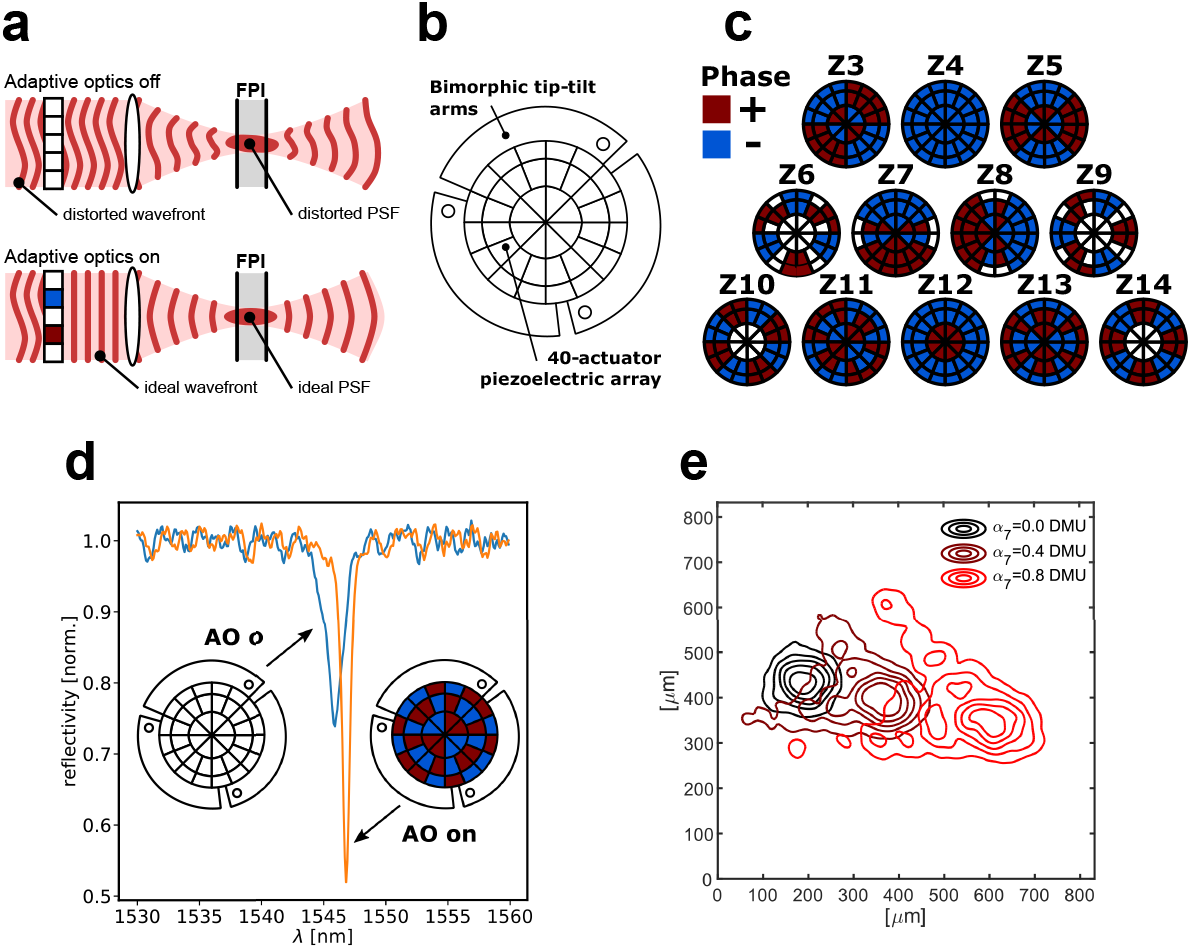
**a)** A schematic illustrating the role of adaptive optics in optimizing the light coupling into the FPI cavity. **b)** Schematic of the deformable mirror used to correct the aberrated wavefronts via modulation of its 40 individual segments. **c)** Basic patterns displayed on the mirror recapitulating low order Zernike aberrations (Z3-Z14, top left to bottom right). **d)** Exemplary FP transfer function at the same sensor position with and without adaptive optics correction **e)** Interrogation beam shape and positions measured on the sensor surface depend on the magnitude of the aberration displayed by the deformable mirror, here shown for coma aberration (Z7).

## 2. Materials and Methods

### 2.1. Experimental AO implementation

We chose an indirect wavefront sensing approach in which individual Zernike modes are applied to the interrogation beam and their respective amplitudes optimized [8, 9]. To impart well controlled phase shifts onto the beam we utilized a Deformable Mirror (DM, DMP40/M-P01, Thorlab, 40 active elements), which was factory calibrated and enabled direct application of Zernike modes Z3-Z14. For most of the considerations in this paper the absolute values of the aberrations are not important and as such the experiments were done using the in-built DM aberration amplitude scale, given as Deformable-Mirror-Units [DMU]. The conversion from DMU to phase units (waves) for a 1550 nm beam is given in **Table S1**.

The implementation of AO hardware on our PAT system is based on an addition of a AO module in front of an optical FP-based PA read-out system (**Figure S1**, see also [6]). The output of the tunable interrogation laser (New Focus, Venturi™ TLB-8800-H-CL) is collimated by a lens (L1, AC254-40-C, f=40 mm) to match the beam size to the diameter (10 mm) of the DM, the beam is then redirected to the setup via a quarter waveplate (*λ/*4) and a polarising beam splitter (PBS). This 90^*o*^ configuration simplifies the alignment of the DM. The surface of the DM is then relayed onto the back focal plane of the scan lens (TSL-1550-15-80, Wavelength Opto-Electronic) by two lens pairs, (L2, AC254-100-C, f=100 mm) - (L3, LE1929-C+#45-805, ef=60 mm) and (L4, LE1929-C+#45-805, ef=60 mm) - (GL, 2x AC254-75-C, ef=37.5 mm).

### 2.2. Photoacoustic imaging and image reconstruction

All *in vivo* mouse experiment procedures were approved by the EMBL Institutional Animal Care and Use Committee (IACUC). A water based gel was placed between the skin and the FPI sensor to provide acoustic coupling. Body temperature of the mice was kept constant during the experiments using an incubation chamber surrounding the FPI. The diameter of the excitation beam (SpitLight DPSS EVO I OPO 100 Hz, InnoLas Laser Gmbh) incident on the skin surface was *≈*1.5 cm with fluence *≈*1 mJ cm^-2^ which is within the safe maximum permissible exposure range for skin [10].

The field-of-view on the FPI sensor was 10 × 10 mm^2^ and scans were acquired from 10,000 positions, with each waveform spanning over *≈*10,000 time points (sampling rate 125 MHz, ATS9440-128M, AlazarTech). The overall tomographic image acquisition time was ≈10 min and was limited by the response time of laser tuning and DM control. The effective acoustic detector size was approximately 80 µm (given by the diameter of the focused interrogation laser beam).

PA images were reconstructed from the raw data using the following steps: The acquired PA signals were interpolated onto a three times denser spatial grid. The speed of sound in the tissue was estimated using a data driven autofocus approach [11]. 3D-images were reconstructed from the interpolated signals using a time-reversal algorithm [12] using the speed of sound estimated in the previous step as a parameter. The image reconstruction was done using an open-source Matlab toolbox (k-Wave [13]).

### 2.3. Quantification of µFPI optical sensitivity

Multiple approaches exist for the quantification of µFPI optical sensitivity from raw ITF data [5, 14]. We chose an approach based on fitting of the Psuedo-Voigt function [14] as this allows for robust and real-time fitting of the µFPI transfer function: 

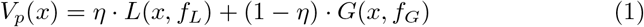

with *L*(*x, f*_*L*_) being the Lorentz function and *f*_*L*_ its FWHM parameter, *G*(*x, f*_*G*_) being a Gaussian function with *f*_*G*_ its FWHM parameter, and *η* is chosen according to ref. [15] as: 

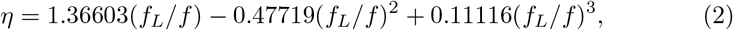

where 

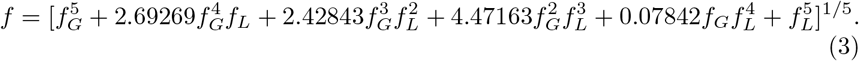

Based on the above fit we calculate the normalised optical sensitivity [6]: 

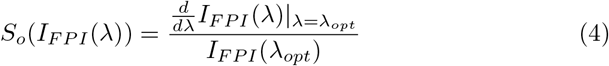

where: 

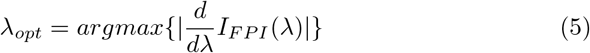

and *I*_*F P I*_ (*λ*) is the transfer function of the interferometer.

## 3. Correction of DM induced focal shifts

Optical aberrations present in laser beams originate from deviations from the Gaussian phase or amplitude profile of the beam and are introduced either through imperfections in the optical elements, their alignment or due to surface inhomogeneities of the FPI surfaces. These deviations can be corrected with the use of active optical elements such as deformable mirrors (DMs, **Figure 1b**) which can be pre-calibrated in the so-called Zernike modes basis which provides a convenient basis for experimental AO correction (**Figure 1c**). Ensuring the phase profile of the laser beam matches the local FPI cavity shape can lead to significant improvements of its optical sensitivity (**Figure 1d**), in line with our earlier theoretical work [6].

One major challenge of using AO correction in photoacoustic tomography (PAT) is the fact that Zernike mode corrections will effectively induce a lateral shift and deformation of the interrogating laser spot on the surface of the FPI dependent on the (Zernike) mode and amplitude (**Figure 1e**). These DM-induced shifts can be significant compared to the spot size for particular aberration modes (**Figure 2a**). This results in two detrimental effects, both of which

**Figure 2:**
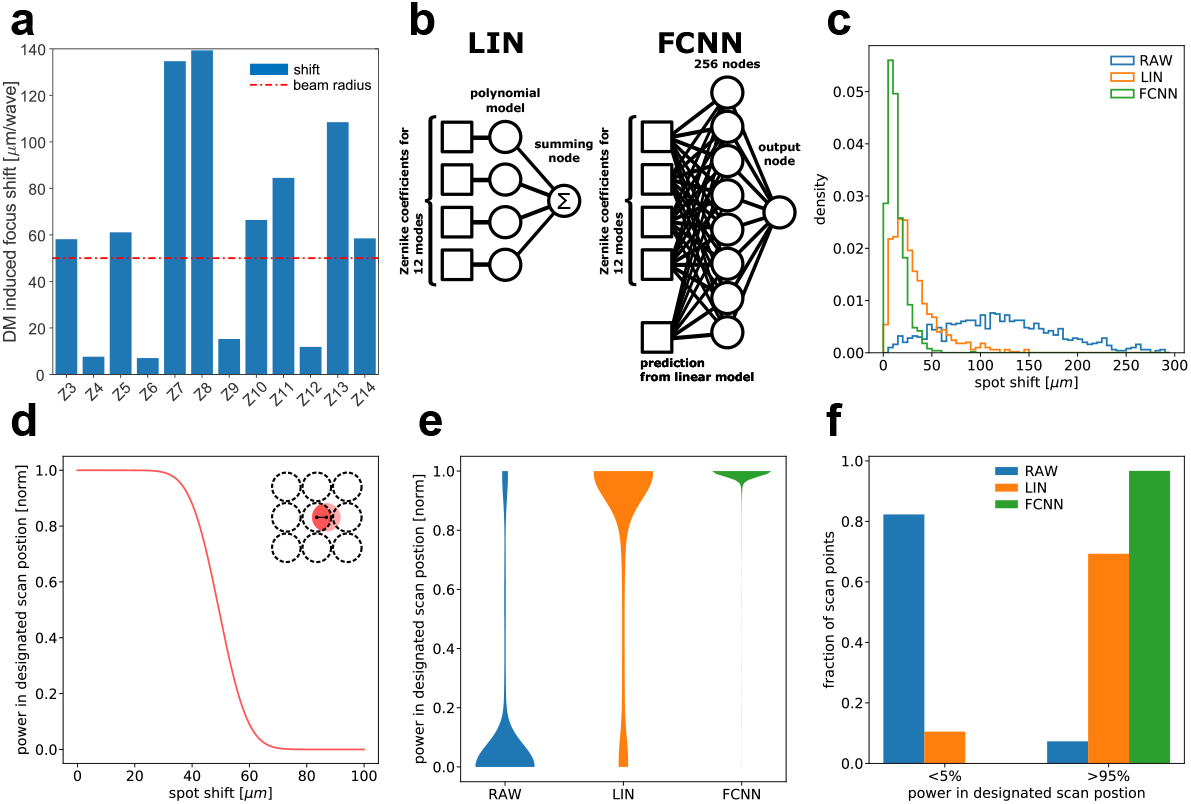
**a)** Experimentally measured DM-induced focus shift for individual Z-modes. Several modes exhibit lateral spot shifts that are larger than the beam radius. **b)** Left: The linear model for shift prediction represented in a network convention. Right: The schematic of the fully connected neural network (FCNN) consisting of a single layer with 256 nodes. Here, the input consists of the 12 Zernike coefficients with the prediction from the linear model serving as a prior. **c)** Histogram of experimentally measured focal spot shifts for a randomly generated sets of aberrations 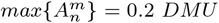 . Here, using the linear model (LIN) as well as our neural network (FCNN) leads to significantly reduced spot shifts compared to the uncorrected case (RAW). **d)** Power contained within the designated scan position on the FPI dependent on the focal spot shift (see **Equation 8**). **e)** Violin plots visualising the loss of power from designated scan position (from **a**) induced by the DM shifts for the uncorrected case (RAW), and after correction based on LIN and FCNN models. **e)** Binning of data from **e** shows drastic differences in predicted performance between the corrected (LIN, FCNN) and uncorrected (RAW) use of the DM.

significantly reduce image quality in PAT: **(1)** Lateral spot shifts effectively deform the scan grid which directly affects the 3D image reconstruction quality as most current image reconstruction methods assume a uniformly spaced grid[13]. **(2)** Since cavity imperfections and thus aberrations are also spatially different, this creates an undesired feedback into the AO optimization routine that can prevent the algorithm to converge to the optimal wavefront correction. This requires careful characterization as well as compensation of these DM-induced focal shifts before and/or during the experiments. For this we developed a novel hardware-based approach, as their effects can not be negated in post-acquisition data processing.

### 3.1. Predicting the spot drift

As real time measurement of the spot drift during AO correction is challenging and unpractical in realistic imaging conditions, we decided to built a model to predict the shift based on the Zernike coefficient of the applied correction. The model exploits the fact that, since Zernike polynomials are orthogonal, their effect on the focal spot shift should be largely independent from each other (linear model (LIN), **Figure 2b**): 

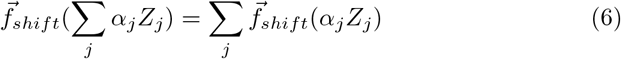

This model can be fairly straightforwardly realised experimentally by measuring the shift induced by single Zernike aberrations of varying amplitudes and fitting low order polynomials (n=3). Doing so yields a predictive model: 

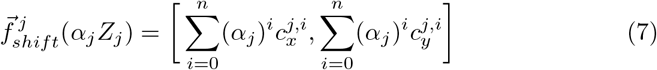

where *α*_*j*_ is the amplitude of the applied correction for mode *Z*_*j*_ and 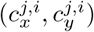 are the fit parameters.

Compared to the uncorrected case (RAW), this linear model (LIN) drastically reduces the overall spot shift (**Figure 2c**). However, the shift itself is not necessarily a good indicator of the actual functional improvement gained by applying the correction. As for our application proper spatial sampling of the photoacoustic field is key, we defined a quality metric based on the optical power fraction contained in the desired scan position for different shift values: 

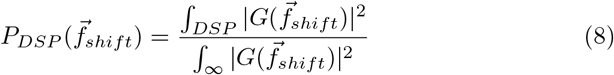

where, *DSP* is the Desired Scan Position and *G* is a Gaussian beam. E.g., for a 100 *µm* beam diameter and a 100 *µm* step size grid this results in a sigmoid behaviour with a rapid loss of contained power around 50 *µm* (**Figure 2d**).

This metric allows us to quantitatively evaluate the effects of DM induced drift and to compare different correction models (**Figure 2e**). We find that for weaker aberrations (i.e. *DMU <* 0.1) the linear model is sufficient to correct the main effects of the drift (**Figure S2e, f**), however for stronger aberrations (i.e. *DMU >* 0.1) we observe a large fraction of points exhibiting signification power loss.

We attribute the shortcomings of the linear algorithm to the fact that the underlying assumption of Z-mode independence is not met for stronger aberrations, presumably because of imperfections in the functioning of the mirror. To address this limitation, we developed a model based on fully connected neural networks (FCNNs) that can account for mode interactions in an agnostic, data driven way (**Figure 2b**). Despite its simplicity, this FCNN can efficiently learn cross-interactions between Zernike modes from a moderate data size of 5000 random aberrations when provided with a prior in the form of the linear model prediction. Therefore, the FCNN outperforms the linear model for both strong and weak aberrations (**Figure S2a, d**) and even in case of strong aberrations limits power losses of the desired scan grid to below 5% in over 96% of scan positions (**Figure 2f**).

## 4. Indirect wavefront sensing with iterative refinement

Having developed an active shift correction scheme, we explored the use of our active adaptive optics routines for enhanced acoustic sensing with FPIs. We chose an *indirect* wavefront sensing approach in which we directly optimized the sensitivity of the system. In contrast, using a *direct* wavefront sensing approach [8, 9] would not only be challenging in its technical implementation, but would also require a complex modeling approach to relate the observed wavefront distortions to changes in sensitivity because of the non-trivial interactions between beam and cavity aberrations in FP interferometers [6].

Aberrations present in our FPI system predominantly come from two sources: Beam aberrations (that are assumed to be **weakly** varying between scan positions on the FPI) and cavity aberrations (that are assumed to be **strongly** varying between scan positions on the FPI). Therefore the overall aberrations are expected to vary significantly between scan positions on the FPI. This presents an experimental challenge since characterising the entire FPI field-of-view in a point-by-point manner would require impractically long optimization time. This is because for iterative AO where e.g. 12 Zernike modes are optimised, each iteration takes *≈*30*s* per scan position which would translate to*≈* 80 hours for the full field of view (10^4^ scan positions). Practically this would extend even further as multiple iterations (2-3) over the same modes are required to compensate for cross coupling between Zernike modes.

However, since aberrations do partially come from beam aberrations which are spatially varying only weakly, a certain extent of spatial correlation is expected. Capitalising on this, we developed a sub-sampling scheme that drastically reduces the overall characterisation time while still providing significant sensitivity improvements over the whole field-of-view. We hypothesised that the same AO correction can still cause sensitivity improvement in the neighbourhood of the point on the FPI on which the characterisation was performed (reminiscent of the ‘isoplanatic patch’ concept in AO microscopy). We therefore divided the whole 99×99 point scan grid into 9×9 coarse regions, and applied the measured AO correction of the center point to the entire coarse region. This approach reduces the time required to characterise the interferometer by two orders of magnitude, i.e. to within one hour. We observed a significant improvement of the sensitivity (**Figure 3a,b**) for almost all FPI positions as is evidenced by pair-wise plotting of the sensitivity values for all AO versus AO ‘off’ scan positions (**Figure 3c**). Here, the magnitude of the improvement can be inferred as the vertical distance from the diagonal and shows that a vast majority of the scan point display significant enhancement in sensitivity.

**Figure 3:**
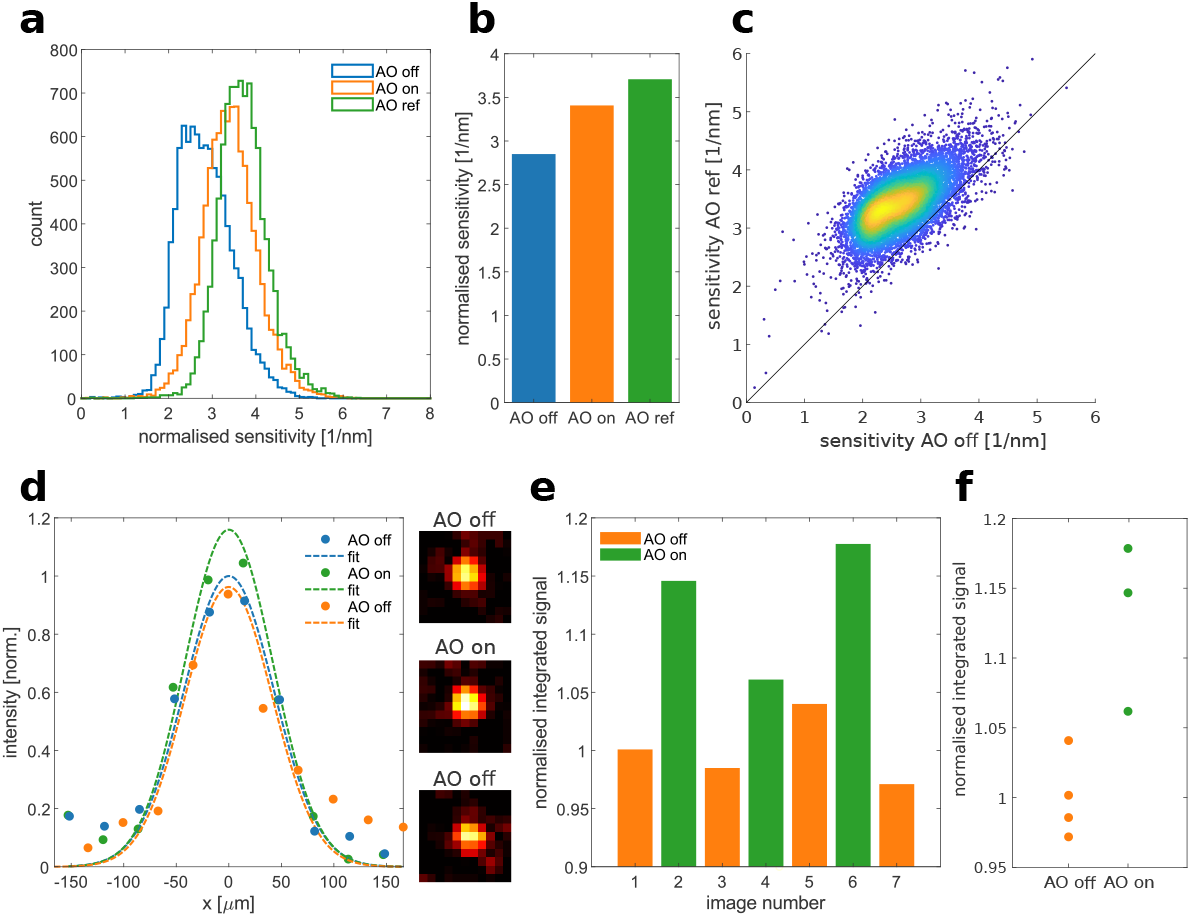
**a)** Histograms of the FP interferometer sensitivity between AO ‘off’ and AO ‘on’ conditions. AO ref refers to the ‘refined’ AO case. **b)** Mean sensitivity of the FP interferometer between AO ‘off’ and AO ‘on’ conditions. **c)** Pointwise correlation between the AO ‘off’ and AO ‘ref’ sensitivity for all the scan positions on the FPI. **d)** Experimentally measured cross-sections of diffraction limited beads taken with the photoacoustic system with AO ‘off’, AO ‘on’ and AO ‘off’ again, respectively, showing improvement for the AO ‘on’ condition. Insets show images from which the cross-sections were taken. **e)** Integrated signal level for consecutive bead images taken while alternating between AO ‘off’ and AO ‘on’ conditions, showing consistently higher signal levels for AO ‘on’. **f)** Dot plot for data shown in **e)**. The images of the beads were corrected for bleaching prior to quantification by fitting an exponential decay curve to the AO ‘off’ conditions and normalising the intensities.

Because the surface of the interferometer is largely subsampled for calculating the AO correction we next investigated to which extent the AO improvement could be refined through denser sampling. This can be achieved either by using a denser grid in the initial characterisation step or by refining the corrections during a second round. We explored the latter option by randomly refining points on the FPI scan grid which were left out during the first characterization and compared the AO-improvement achieved in a 9×9 region centered around this point with the previous AO-improvement, updating the correction where it yielded an improvement in sensitivity. This refinement approach, termed AO ‘ref’, allowed us to further enhance the overall sensitivity of the interferometer in an iterative fashion (**Figure S3**) achieving an improvement over the standard AO ‘on’ condition (**Figure 3a,b**). This iterative refinement leads to an overall increase in (mean) sensor sensitivity with additionally refined scan positions, which can be described by a saturating function (**Supplementary Section S1,Figure S3**). This allows to predict the sensitivity improvement with the number of additionally refined scan positions and thus to define a stopping condition. In this study we chose to refine an additional 500 points (5% of the scan grid) which took *∼* 8h. We did not refine more points as significant saturation could already be observed (**Figure S3**).

The improvement in sensitivity due to AO comes from an increased slope of the ITF, however it also affects the visibility and shifts the overall signal level. This DC signal change also entails an actual shift of the working point of the photodetector, which goes in hand with DC related noise sources such as the relative intensity noise (RIN) and shot noise, therefore affecting the overall performance of the system. To account for this effect we developed a simple ap-proach which allowed to correct the input power of the PA system to equalize the working point in both conditions (**Figure S4**, for details see **Supplementary Section S2**). This is important to ensure that any improvements in coupling of light into the FPI due to the AO actually translate into improved overall sensitivity of PAT imaging.

## 5. Adaptive-optics enhanced photoacoustic imaging

To validate the potential of our approach for improving PAT we performed imaging experiments using resolution phantoms made of 10 µm sized dye loaded beads (1010KB, Degradex^®^) embedded in 1% agarose and quantified the improvement in signal level when using AO enhanced cavity coupling (**Figure 3d**). We observed that the peak intensity of the reconstructed bead images increases when AO corrections are applied. To further ascertain that the observed improvement indeed came from applying adaptive optics we performed sequential AO ‘on’ and ‘off’ imaging and observed a reproducible pattern showing clear improvement whenever AO was turned on (**Figure 3e,f**).

We then proceeded to apply our approach on biological samples by performing *in vivo* label free imaging of hemoglobin in the abdominal area of anaesthetised mice. Using our developed AO correction scheme described above we were able to improve PA signal levels under realistic imaging conditions (**Figure 4a**). In particular, AO led to a strong enhancement of signal level in the recorded PA frequency band of 1-20 MHz (**Figure 4b**), as quantified by calculating the power spectrum of the recorded PA signals. Integrating the power spectrum over this frequency bandwidth shows a significant 3-fold improvement in signal level (**Figure 4c**). Due to the spatial heterogeneity of the FPI sensor used in PAT we expect the magnitude of this improvement to also vary spatially, causing stronger local enhancement of image quality. By comparing sequentially taken PA images (**Figure 4d-f, top row**) we observe that, while global quality metrics only show a moderate improvement in image quality (**Figure 4g, top**), locally we can observe much larger improvements (**Figure 4d-g, bottom row**). This is however expected given the spatial variation in AO improvement measured across the sensor.

**Figure 4:**
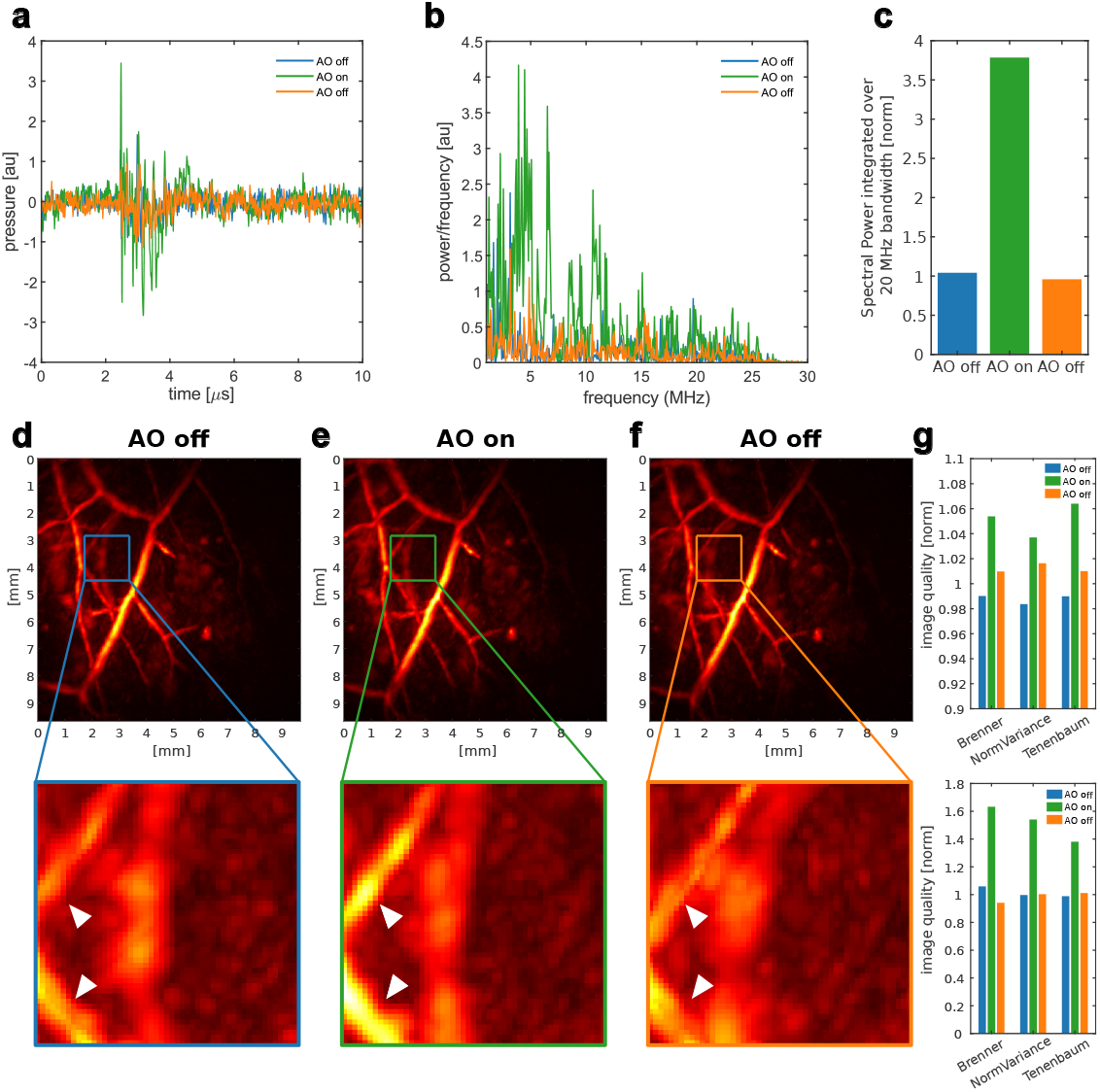
**a)** Raw PA time traces from an in-vivo imaging experiment comparing signal level between AO ‘off’ and AO ‘on’ conditions. **b)** Power spectrum from time traces in panel **(a)** showing frequency enhancement in the 1 - 30 MHz band. **c)** Quantification of the overall increase in signal level in the 1 - 30 MHz band by integrating the power spectrum from panel **(b). d-f)** Maximum intensity projections of photoacoustic images of the abdominal part of the mouse taken at 600 nm showing label-free imaging of vasculature (top), as well as zoom-ins to regions that show especially good image quality improvement (bottom, see arrows). **g)** Commonly used image quality metrics calculated for images of the same area taken with AO ‘off’, AO ‘on’ and AO ‘off’ again, for both the whole images (top) and the zoomed in regions (bottom), respectively.

## 6. Discussion

In this work we have developed an experimental paradigm that utilizes adaptive optics to enhance the sensitivity in all-optical photoacoustic tomography based on Fabry-Pérot interferometers and that allows to translate these improvements to practical imaging conditions. This was achieved by a careful characterization of the influence of active wavefront shaping on the position and shape of the interrogating beam on the sensor, and by actively adjusting the interrogating beam power on the photodetectors to ensure optimal SNR. Furthermore, we presented a subsampling approach that allows to characterize and iteratively refine the AO corrections on a practical time-scale. Altogether, our AO approach enabled us to enhance photoacoustic signal detection by up to 3.5-fold, and to improve image quality in in-vivo experiments.

We want to highlight that the use of AO in our PAT system differs significantly from its traditional use in (light) microscopy. In our case, AO is utilized to improve the sensitivity of the photoacoustic readout. This, however, is contrary to the typical application of AO in microscopy where it is used to yield improvement in spatial resolution. Recently, computational PA wavefront correction techniques were developed that allow for adaptive photoacoustic tomography in this traditional meaning [16].

While we report significant improvements in both sensitivity and image quality, those gains do not restore the theoretically possible maximum sensitivity of FPI sensors. We have previously discussed potential (theoretical) reasons why a full recovery of sensitivity via AO is not attainable in a realistic systems [6]. In essence, cavity aberrations amplify the optical aberrations present in the interrogating beam while traveling within the FPI cavity. Fully correcting these would require controlling the phase of the beam simultaneously at multiple focal planes (corresponding to the multiple reflections inside the FPI) which is not possible as the phase evolution is determined by the wave equation. This limits AO to only partially correcting the effects of cavity aberrations. Here, the scope of our study was limited to lower order aberrations due to the low resolution of the DM. Utilizing active wavefront shaping devices with more active elements such as spatial light modulators (SLMs) could allow to correct for higher order aberrations and thus to further improve the sensitivity of the FPI. Eventually, future gains in overall sensitivity will likely also require advanced manufacturing techniques that can actively alter the cavity structure locally [17, 18] and/or can ensure high thermal stability of the FPI cavity.

Finally, we note that our work also indirectly explores the possibilities of wavefront shaping. Because of the presence of cavity aberrations, the best beam profile to interrogate the FPI might not necessarily be a Gaussian beam.

As a result, the iterative indirect wavefront sensing approach used in this paper will possibly generate non-Gaussian beams that might better suit particular local cavity shapes of the FPI. In this sense, our approach goes beyond pure aberration correction. Unfortunately, as the DM employed in our work has only limited degrees of freedom it was not possible to explore more complex beam profiles that are far from the fundamental Gaussian beam. Here again, the use of SLMs possessing thousands of pixels will allow to further explore this regime. Among others, recent work has shown the use of non-Gaussian beams to increase FP measurement sensitivity, e.g. by utilizing *LG*_33_ modes in LIGO detectors [19], and Bessel beams have been employed in FP micro-cavities [20] to enhance the sensitivity of FPI pressure detection.

## Acknowledgments

We would like to thank the EMBL Heidelberg mechanical and electronic workshop as the animal facilities for help and support. We further thank Florian Mathies, Johannes Zimmermann, and Gerardo Hernandez-Sosa from InnovationLab Heidelberg as well as Karl-Phillip Strunk and Jana Zaumseil from the Centre for Advanced Materials, Heidelberg University, for help with manufacturing of the Fabry–Pérot interferometers used in this work. We would also like to thank Carlo Bevilacqua and Nikita Kaydanov for critically reading the manuscript and providing feedback as well as Lina Streich for helpful discussions on adaptive optics and the operation of deformable mirrors and Martin Jechlinger for support with the animal experiments. This work was supported by the European Molecular Biology Laboratory.

