## Supplementary Information for "Adaptive optics enhanced all-optical photoacoustic tomography"

### Adaptive optics enhanced all-optical photoacoustic tomography: Supplementary Information

This document provides supplementary information on the main manuscript "Adaptive optics enhanced all-optical photoacoustic tomography". It contains the discussion on modeling of the adaptive optic refinement and optimal working point of the photodiode as well supplementary tables and figures.

#### S1 Modeling the adaptive optic refinement

As the iterative AO improvement described in **Section 4** is a saturating process the improvement of mean sensor sensitivity with additionally refined sensor positions can be nicely approximated by the following function inspired by the Michaelis-Menten equation:

$$S_o(n) = S_{max}\frac{n}{k_s + n} + S^0 \tag{S1}$$

where, $S_{max}$ is the maximum sensitivity improvement, $k_s$ is the iteration number at which we reach half of $S_{max}$, $S^0$ is the starting sensitivity and $n$ is the number of additionally characterised points on the FPI. This allows us to quantify the sensitivity improvement versus time trade-off and choose and optimal stop condition.

#### S2 Optimal working point of the photodiode

We developed a simple model to account for the changes in the photodiode working point. Due to spatial heterogeneity of the FPI cavity the intereferometer will display a distribution of normalised sensitivities ($S_i$) and a corresponding distribution of working points ($p_i$). This is due to different degrees of reflectivity at the bias wavelength which leads to varying levels of DC power on the photodetector. The effective sensitivity at each point is linearly dependent on the incident power provided that the power level does not exceed the saturation limit of the detector ($P_{saturation}$). The DC power level at each point can be defined as $P_i = Pp_i$ where P is the output power of the interrogation laser and $p_i$ is the working point defined as the fraction of the laser power incident onto the photodetector for a particular position on the FPI when tuned to the bias wavelength. Because of this the effective normalised sensitivity at each point will depend on the photodetector being saturated or not:

$$S(P_i) = \begin{cases} 0 & \text{if } P_i \geq P_{saturation} \\ S_i & \text{if } P_i < P_{saturation} \end{cases} \tag{S2}$$

Taking this into consideration we can write the expression for the effective signal-to-noise ratio (SNR) in the reconstructed image. For simplification, we assume that each point on the surface of the FPI detects signals from the entire imaging volume which are coherently summed in the reconstruction process, therefore the sensitivity at each point of the reconstructed volume equals:

$$S \sim \sum_{i=0}^{n} S(P_i)P_i \tag{S3}$$

[1]Cell Biology and Biophysics Unit, European Molecular Biology Laboratory, Heidelberg, Germany
[2]Developmental Biology Unit, European Molecular Biology Laboratory, Heidelberg, Germany
[3]Epigenetics and Neurobiology Unit, European Molecular Biology Laboratory, Monterotondo, Italy
[4]Molecular Medicine Partnership Unit, European Molecular Biology Laboratory, Heidelberg, Germany
[5]Collaboration for joint PhD degree between EMBL and Heidelberg University, Faculty of Biosciences, Germany

Similarly we assume that the system is shot noise limited and the noise at each point adds incoherently:

$$N \sim \sqrt{\sum_{i=0}^{n} P_i} \tag{S4}$$

Therefore the expression for the effective SNR becomes:

$$SNR_{eff} = \frac{S}{N} \sim \frac{\sum_{i=0}^{n} S(P_i)P_i}{\sqrt{\sum_{i=0}^{n} P_i}} \tag{S5}$$

The output power of the interrogation laser (P) is a free parameter which allows us to optimise the performance of the setup for a give $p_i$ and $S_i$ distribution which can be measured experimentally.

#### S3 Supplementary Tables

| Zernike mode | Aberration | Scaling [wave/DMU] |
|---|---|---|
| Z3 | Oblique astigmatism | 8.77 |
| Z4 | Defocus | 8.39 |
| Z5 | Vertical astigmatism | 8.77 |
| Z6 | Vertical trefoil | 3.10 |
| Z7 | Vertical coma | 3.23 |
| Z8 | Horizontal coma | 3.23 |
| Z9 | Oblique trefoil | 3.10 |
| Z10 | Oblique quadrafoil | 2.71 |
| Z11 | Oblique secondary astigmatism | 1.42 |
| Z12 | Primary spherical | 1.29 |
| Z13 | Vertical secondary astigmatism | 1.42 |
| Z14 | Vertical quadrafoil | 2.71 |

Table S1: Scaling factors to convert from Deformable Mirror Units [DMU] to waves for a 1550 nm laser beam

#### S4 Supplementary Figures

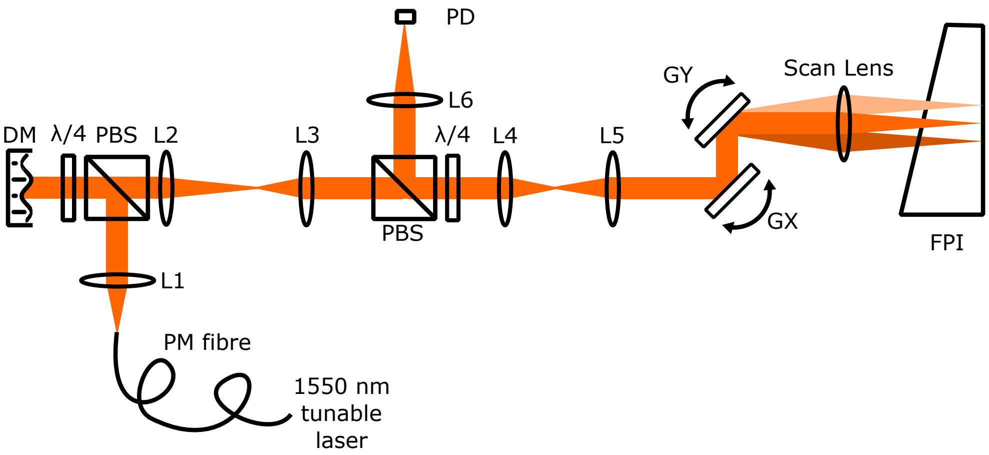


Figure S1: Setup scheme. **DM** - deformable mirror; **LX** - lens X; **λ/4** - quarter waveplate; **PBS** - polarising beam splitter; **PD** - photodiode; **GX,GY** - galvanometric mirrors X and Y, **FPI** - Fabry-Pérot interferometer

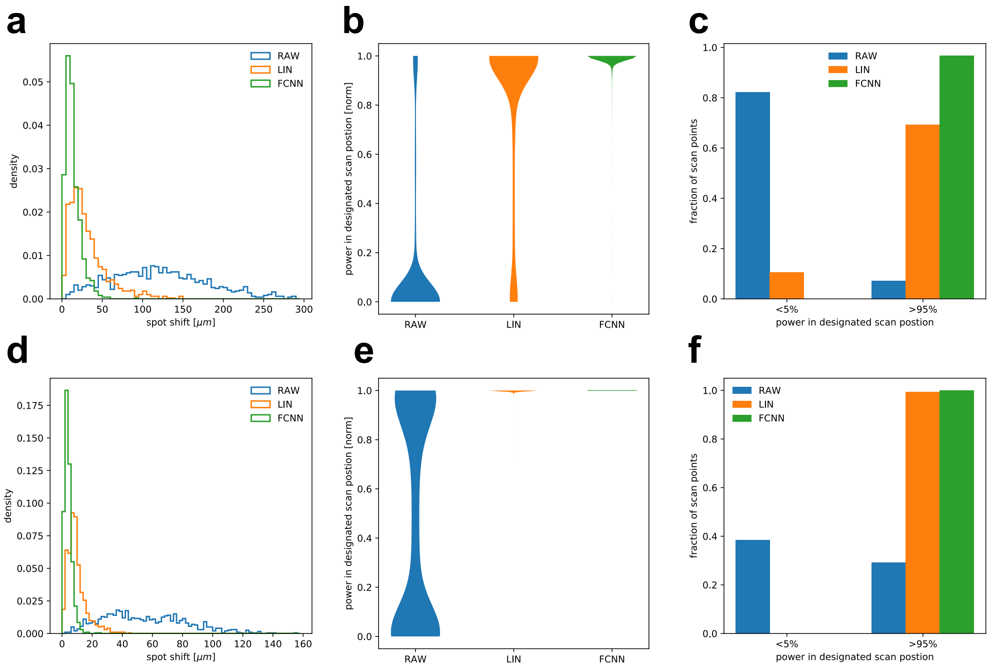


Figure S2: Comparison of focal shift corrections. Panels **a,b,c** are replotted from **Figure 2** for better comparison ($max\{A_n^m\} = 0.2\ DMU$). Panels **d,e,f** are analogs of panels **a,b,c** respectively, but gathered for aberrations where $max\{A_n^m\} = 0.1\ DMU$.

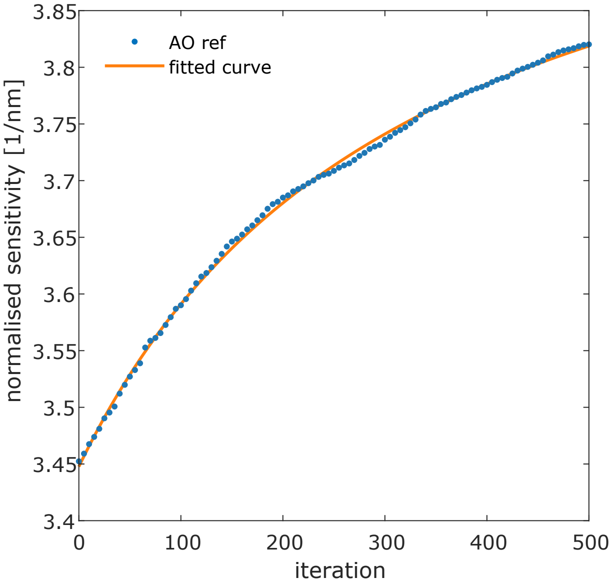


Figure S3: Iterative sensitivity improvement over the number of additionally characterised points using AO refinement and the fit with a saturating function (**Equation S1**).

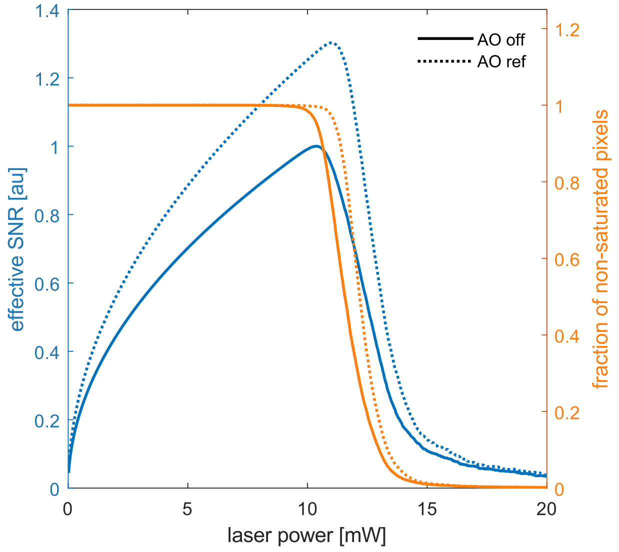


Figure S4: Effective SNR as a function of interrogation laser power incident on the photodiode, comparing between the AO 'off' and AO 'ref' conditions as described by our model.
